# Enhanced production of natural shiga toxin by overexpressing A subunit of Stx2e in Stx2e-producing *Escherichia coli* isolated in South Korea

**DOI:** 10.1101/2022.07.30.502126

**Authors:** Byoung Joo Seo, Jeong Hee Yu, Seung Chai Kim, Yeong Ju Yu, Byung Yong Park, Jung Hee Park, Won Il Kim, Jin Hur

## Abstract

This study explored the optimal culture conditions for maximizing shiga toxin production in Stx2e-producing *Escherichia coli* (STEC) 150229, isolated from porcine edema disease (ED), with the goal of preparing a Stx2e toxoid vaccine candidate. High cytotoxicity was observed for this strain [tissue culture cytotoxic dose 50% (10^4^ TCCD_50_/100 µl)] from 48 h after incubation. Stx2e was overexpressed by transforming pStx2e A into STEC 150229, resulting in the production of recombinant Stx2e A/B complex combined with intrinsic Stx2e B. The enhanced production of Stx2e was evaluated based on the level of cytotoxicity against Vero cells. The highest cytotoxicity (10^5^ TCCD_50_/100 µl) was observed with the samples of recombinant Stx2e A/B complex eluted with 500 mM imidazole at 48 h of incubation. In conclusion, the recombinant Stx2e A protein forms an active protein complex with the intrinsic Stx2e B component from STEC 150229, producing high levels of shiga toxin.

## Introduction

In pigs and wild animals, edema disease (ED) is primarily caused by Stx2e-producing *Escherichia coli* [1, 2]. In humans, STEC also causes hemorrhagic colitis and/or hemolytic-uremic syndrome [3]. Pigs infected with STEC, which is associated with high doses of Stx2e toxin, can exhibit edema at various sites, most notably the colon, stomach, small intestine, eyelids, and brain. Evidence of damage to the vascular endothelium resulting from such edema includes ataxia, recumbence, convulsions, and paralysis [4], which ultimately result in a high mortality rate in Stx2e-producing *E. coli* -infected pigs [5, 6].

Shiga toxin-producing *E. coli* (STEC) produces two major toxin types, Stx1 and Stx2, which are functionally and structurally related to shiga toxin produced from *Shigella dysenteriae* [7]. Even though there is greater than 50% genetic similarity between Stx1 and Stx2 toxins [8, 9], they differ in their structural and immunological characteristics [10]. The animal-related variants of STX2 in STEC, which are not found in humans, are known as Stx2e [11], Stx2f [12], and Stx2g [13]. One of these, Stx2e as a component inducing enterotoxemia of pigs, is mainly found in ED-affected pigs. Structurally, Stx2e toxin exhibits a heterohexameric complex (1A5B type) in which a single A subunit is associated with a homopentamer of B subunits [3, 14, 15]. The heterohexameric complex consists of the A subunit, which plays a catalytic role, and five B subunits acting as glycolipid receptors and in the binding of globotriaosylceramide (Gb_3_) or globotetraosylceramide (Gb_4_), on intestinal epithelial cells [16-18]. In more detail, the A subunit is composed of A1 and A2 fragments, which are tethered by a disulfide bridge between cysteines 241 and 260. The A1 fragment contains an active site that mediates the cytotoxic effects. In contrast, the A2 fragment is located at the central hole of pentameric B subunit. The A subunit exerts N-glycosidase activity suspending protein synthesis within the targeted cells [15]. In this study, a cytotoxicity assay was used to demonstrate the toxin production of STEC isolates from a field case of porcine ED, and enhancement of toxin production was also indicated by overexpressing recombinant N-His Stx2e A/B complex with intrinsic Stx2e B in STEC 150229 (recombinant Stx2e A/B complex).

## Materials and methods

### Ethics Statement

This study was performed in strict accordance with the recommendations of the Guide for the Care and Use of Laboratory Animals of the National Institutes of Health. All animal experimental protocol was approved by the Jeonbuk National University Institutional Animal Care and Use Committee (Approval Number: JBNU 2021-0136) and performed according to accepted veterinary standards set by the Jeonbuk National University animal care center. Mice were euthanized by CO_2_ inhalation, as specified by the Jeonbuk National University Institutional Animal Care and Use Committee guidelines.

### Shiga toxin producing *E. coli* strain

At Chonbuk National University Diagnostic Center, edema disease-causing STEC 150229 was isolated from the feces of pigs from a pig farm in South Korea diagnosed with this disease (CBNU-VDC) [19]. The isolated STEC 150229 strain was routinely grown in LB broth (Affymetrix USB, CA, USA) at 37°C for 48 h. The cell-free supernatants of bacterial strains were prepared by culturing them, centrifugation (10,000 ×g for 5 min) and filtration (0.22-µm membaranefilter; Sartorius Stedim Biotech, Goettingen, Germany).

### Cytopathic effect of Shiga toxin in animal cell line

The cytopathic effect (CPE) was tested in Vero cell line 76 (ATCC CRL-1587). Vero cells were cultivated in Dulbecco’s modified Eagle’s medium (high glucose) (DMEM, #LM001-05; Welgene, Korea) supplemented with 10% fetal bovine serum, 1% L-glutamine 200 mM (100×) (#25030-081; Gibco, NY, USA) and 1% Anti-anti (100×) (#15240-062; Gibco) at 37°C and 5% CO_2_. The Vero cell toxicity assay was performed in 96-well tissue culture plates (#351172; Falcon, NY, USA) using 10-fold dilutions of samples. The plates were analyzed for the presence of CPE after 48 h and tissue culture cytotoxic dose 50% (TCCD_50_)/100 µl was calculated.

### Vector and Toxin gene cloning

The gene of Stx2e A subunit (GenBank ID: CP024997.2) was amplified with specific primer sets and inserted into the modified expression vector with *Nco*I and *Not*I restriction enzyme (Fig. 2, pStx2eA). In order to modify an expression vector, the pelB gene, secretion signal sequence, was connected to the pET30a (New England Biolabs Inc., MA, USA) vector with *Nde*I and *Nco*I restriction site. A six-His tag was fused to the N-terminus of the Stx2e A gene to facilitate protein purification.

To prepare purified Stx2e protein for antibody production in mice, two additional plasmids were constructed; Stx2e A-fragment (amino acid 215-319) was inserted to pTWIN vector with *Nco*I and *Bam*HI restriction enzyme (Fig. 2, pStx2e A-fusion). Stx2e B was inserted to pET30a vector with *Nco*I and *Bam*HI restriction enzyme (Fig. 2, pStx2e B).

### Overexpression and purification of Stx2e and recombinant Stx2e A fusion/Stx2e B (r Stx2e Afrag/B)

The Stx2e A (Fig. 2) was overexpressed in STEC 150229 and *E. coli* BL21 (DE3) cells which cultured on LB broth containing 25 µg/mL of kanamycin (Duchefa, Netherlands). The protein expression was induced with 0.5 mM isopropyl β-D-1-thiogalactopyranoside (IPTG) (Duchefa) at O.D._600_ of 0.4 and the cells incubated for 24 h at 18°C, 170 rpm. The proteins were purified from supernatant with HisTrap FF column (GE Healthcare, PA, USA) and eluted using sequentially increased Imidazole concentration (25, 50, 100, 250 and 500 mM) in 10 mM Tris buffer, pH7.0. The protein obtained from serial elution step was analyzed by 15% SDS-PAGE.

The Stx2e A-fragment and Stx2e B proteins were co-expressed on *E. coli* BL21 (DE3) pLysS cells which cultured on LB broth containing 25 μg/ml of kanamycin, 50 μg/ml of ampicillin (Duchefa) and 34 µg/ml of chloramphenicol (Wako, Japan). The protein expression induced with 0.8 mM IPTG at O.D._600_ of 0.5 and cultures were incubated for 16 h at 18°C, 150 rpm. Overexpressed proteins were purified with FPLC system (GE healthcare) by using HisTrap FF column (buffer A : 10mM Tris-HCl, pH 8.5, 100 mM NaCl and buffer B : 10 mM Tris-HCl, 100 mM NaCl, 500 mM Imidazole, pH8.5).

### Immunization of mice and challenge with recombinant Stx2e toxin

Twenty C57BL/6 female mice were purchased and randomly divided into 2 groups. All groups were intramuscularly primed (IM) at week 6 of age [0 week post prime immunization (WPPI)] and were IM boosted at 2 WPPI. The immunized group mice were immunized with the 100 µg of rStx2e A frag/B. The negative control group mice were injected with sterile phosphate-buffered saline (PBS). To evaluate each the antigen-specific antibody titer, blood samples were collected at 0, 2 and 4 WPPI. All samples were stored at −80°C until use.

## Results

### Optimizing cultivation of Shiga toxin producing *E. coli* strain

To examine the optimal cell count and cytotoxicity levels, STEC 150229 was cultivated for 72 h. It showed its maximal cell count of 10^8^ colony-forming units (CFU)/ml at 24 to 72 h after incubation and a high cytotoxicity level of 10^4^ TCCD_50_/100 µl at 48 and 72 h after incubation (Fig. 1). Based on the results regarding the optimal culture conditions, culture of STEC150229 for 48 h was used in this experiment.

**Fig. 1.**
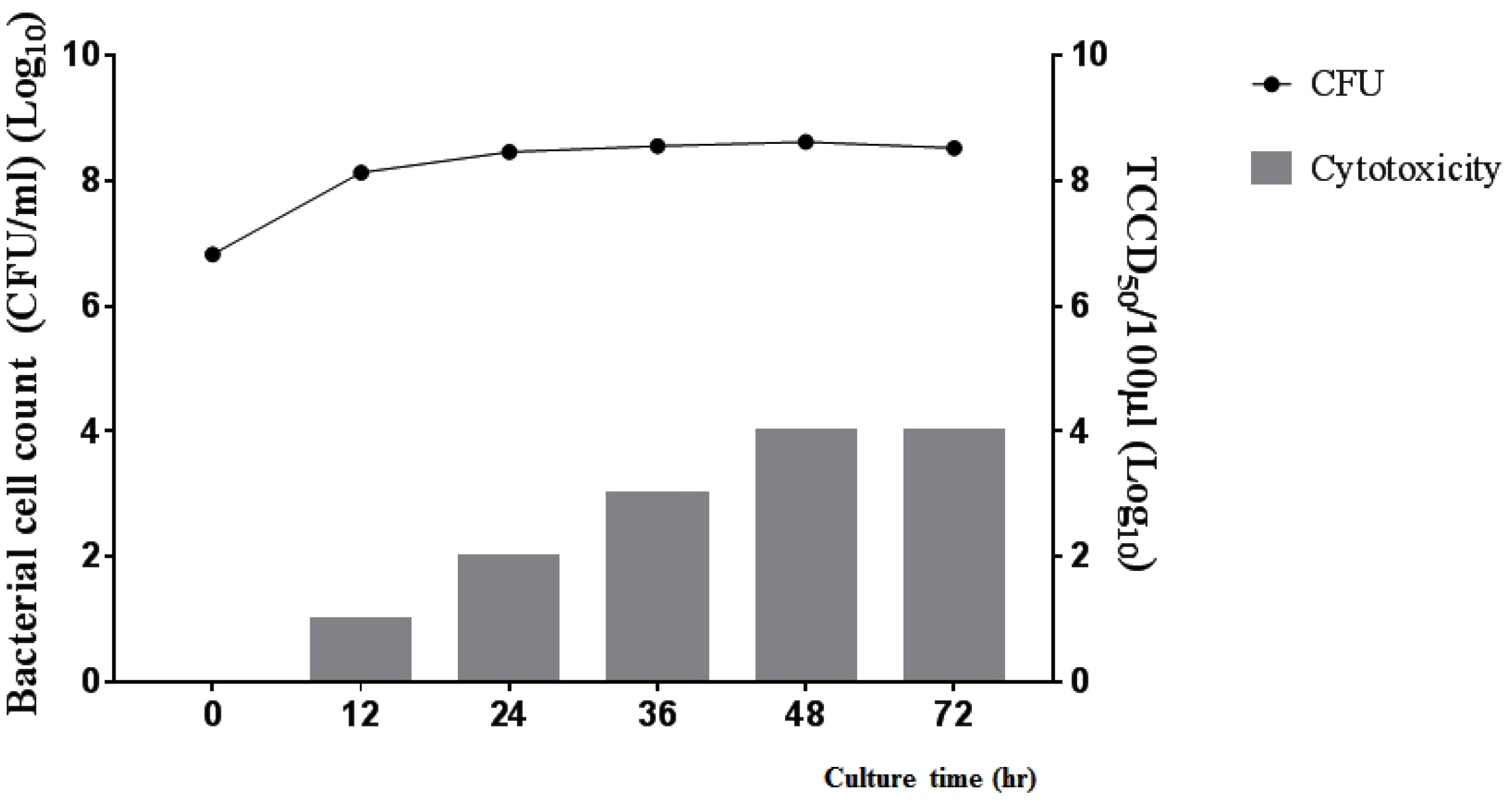
The optimization and cytotoxicity levels of STEC 150229 during incubation.

**Fig. 2.**
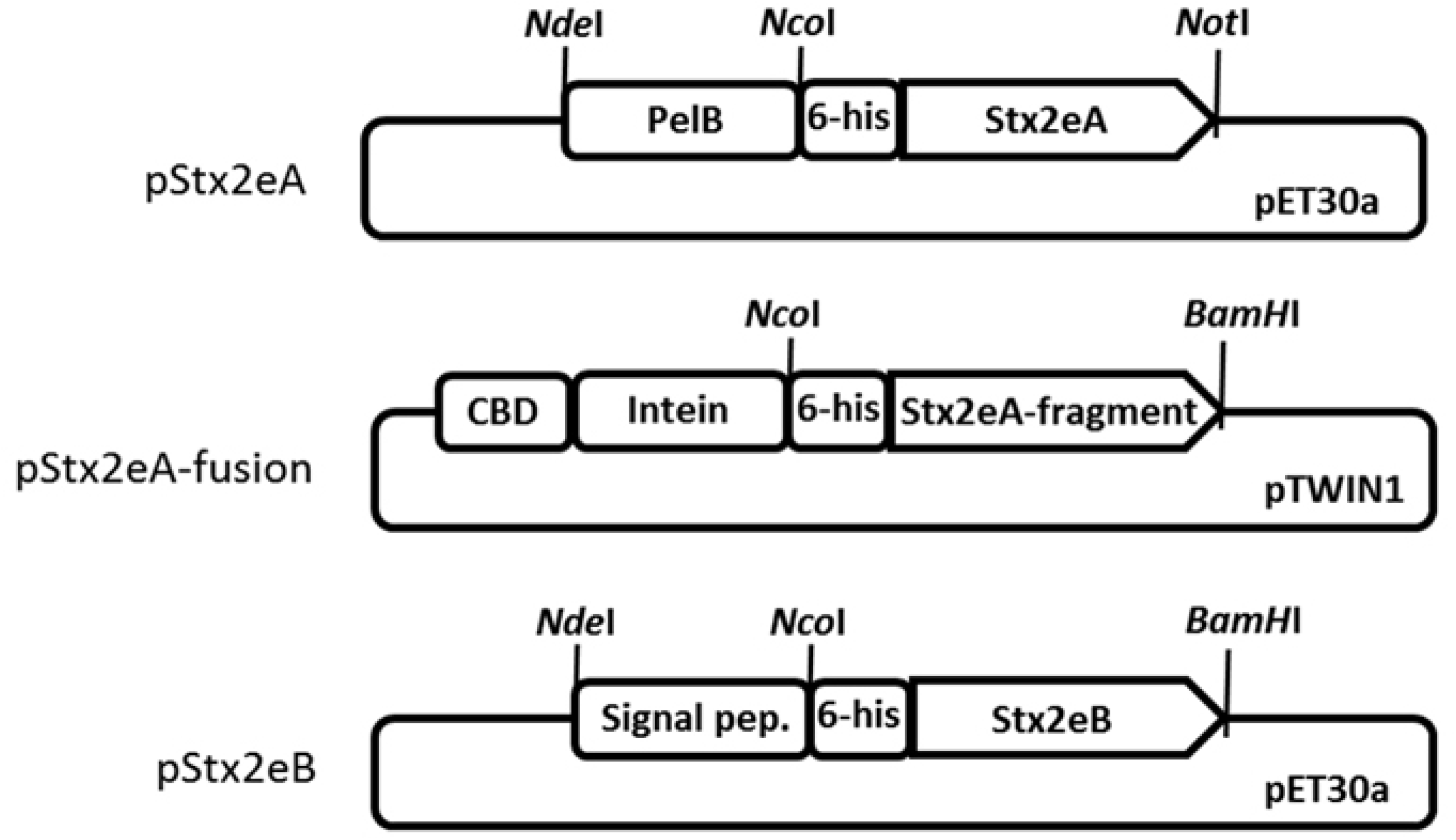
Scheme of the gene constructs of recombinant Stx2e A and B proteins. In pStx2eA plasmid, the pelB signal sequence for periplasmic secretion was connected at the N-terminus of the Stx2eA gene and a 6-histidine tag was inserted in the middle to facilitate purification. In the pStx2eA-fusion plasmid, Stx2eA fragment (a.a. 215–319) was connected to the N-terminus of the intein protein in the pTWIN1 vector. Stx2eB gene was inserted into the pET30a vector, which contained a signal peptide in the N-terminus.

### Production of rStx2e A frag/B complex proteins and immunization of mice

To increase the production of Stx2e complex, we tried to overexpress the recombinant 6-His Stx2e A protein in the STEC 150229 cell. According to our hypothesis, the recombinant protein could establish an enzymatically active complex with the innate toxin proteins. During the purification step, the intrinsic shiga toxin complex was removed using a column at a low imidazole concentration and the recombinant complex was eluted and concentrated (Fig. 3). Our findings showed that a functional Stx2e protein complex was constructed with the recombinant Stx2e A-His and intrinsic Stx2e B in STEC 150229 (rSTEC 150229), the cytotoxicity of which was evaluated in Vero cells.

**Fig. 3.**
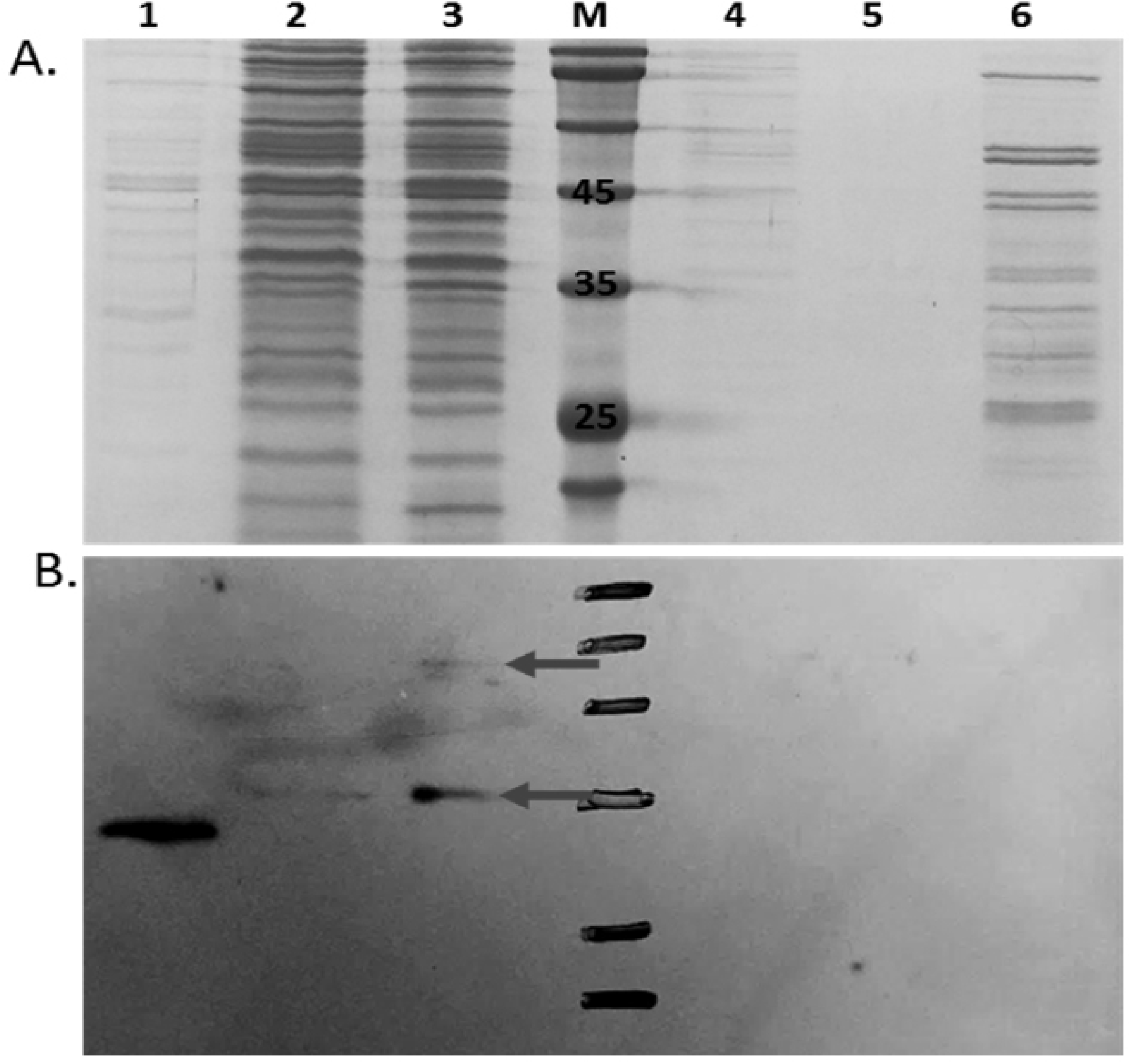
SDS-PAGE (A) and western blotting (B) of recombinant Stx2eA and innate Stx2eB. In lanes 2 and 3, Stx2eA protein was detected (arrow). [Lane M: size marker; Lane 1: *E. coli* BL21 (DE3) cell (negative control); Lane 2: concentrated STEC 150229 culture medium; Lane 3: concentrated Stx2eA-overexpressing STEC 150229 culture medium; Lane 4: flow-through; Lane 5: eluent with 250 mM imidazole; Lane 5: eluent with 500 mM imidazole].

To detect intrinsic and overexpressed Stx2e toxin, an antigen-specific antibody was required. However, the expression level of recombinant Stx2e (6-His Stx2e A and intrinsic Stx2e B) was extremely low, so it was difficult to purify and concentrate it for use as an antigen. Even purified recombinant Stx2e complex concentrated 1000-fold was not detected on western blotting with anti-His antibody (data not shown). In addition, rSTEC 150229 was difficult to inject into mice due to the high production of shiga toxin. Thus, we constructed a new recombinant protein able to produce antibodies and to detect shiga toxin of rSTEC 150229.

Anti-Stx2e antiserum was produced by two injections of rStx2e Afrag/B into C57BL/6 mice. To detect intrinsic and overexpressed recombinant Stx2e toxin, SDS-PAGE (Fig. 3A) and western blotting with antiserum collected from a mouse injected with rStx2e Afrag/B were performed (Fig. 3B). The culture medium harbored secreted innate Stx2e toxin and recombinant toxin complex and purified proteins were concentrated 1000-fold. The estimated molecular weight of Stx2e A is 32 kDa and that of intrinsic Stx2e B is 7.7 kDa (14,15). Based on this estimation, Stx2e A with a size of 35 kDa was detected in STEC 150229 and rSTEC 150229 although the concentration was significantly higher in rSTEC 150229. In addition, Stx2e A/B complex of approximately 55 kDa in size was detected only in rSTEC 150229 (Fig. 3B). The Stx2e A and B proteins purified from rSTEC 150229 cells were not detected in western blotting, even after 1000-fold concentration.

### Cytopathic effect of recombinant Stx2e complex protein

The cytotoxicity of all samples is listed in Table 1. Deletion of the N-terminus of Stx2e A, which is involved in the enzymatic reaction, was associated with the loss of cytotoxicity (Table 1). The cytotoxic activities were not detected in Stx2e Afrag/B protein-overexpressing *E. coli* BL21 (DE3) or in native *E. coli* BL21 (DE3), as a negative control (Table S1).

**Table 1.**
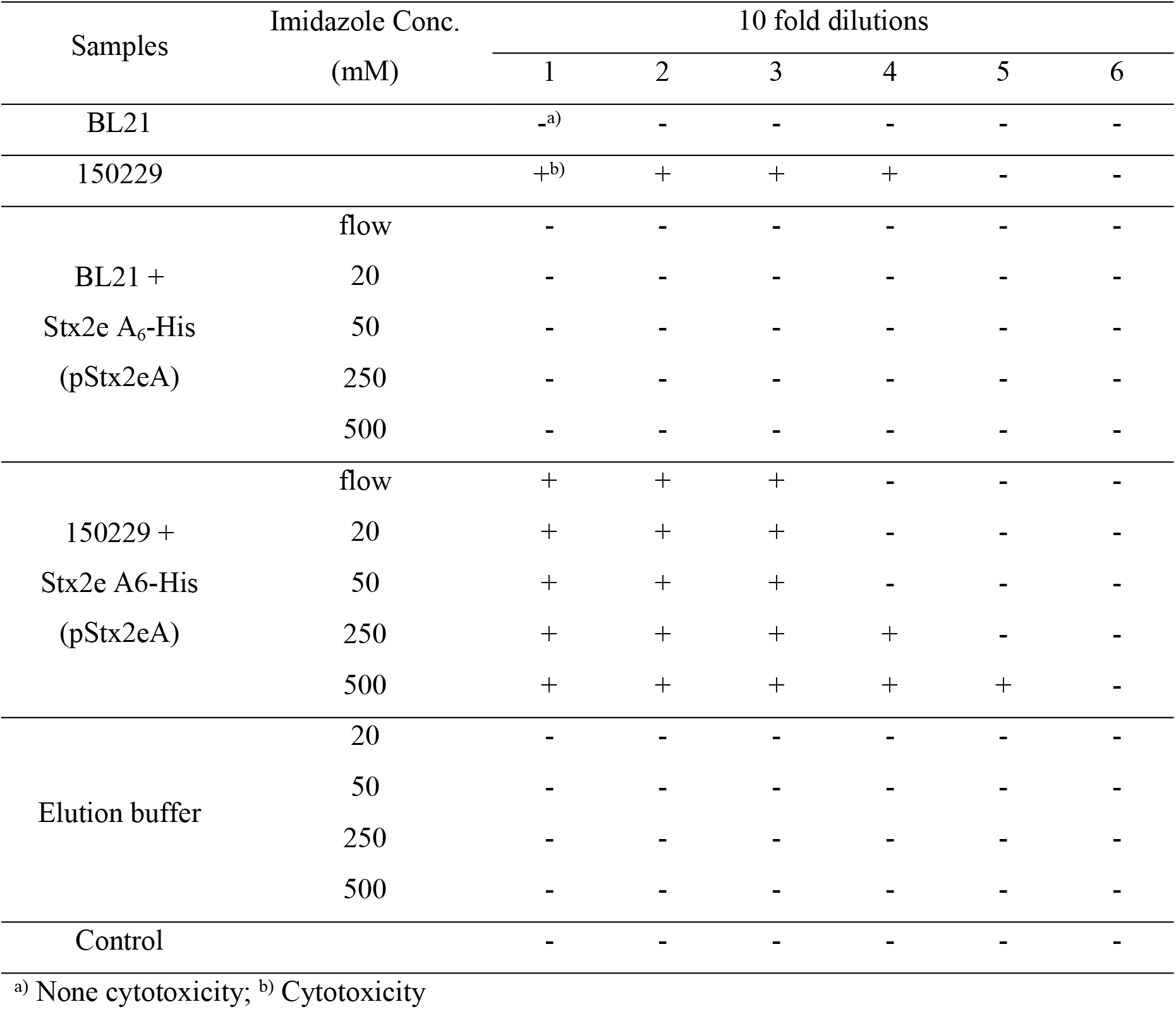
Cytotoxic levels of recombinant Stx2e A_6_-His/B and STEC 150229 in Vero cell

The cytotoxicity of unbound and eluted sample with a low imidazole concentration was caused by the intrinsic toxin complex. The cytotoxic activity of recombinant 6-His Stx2e A/B samples eluted with 500 mM imidazole was significantly higher than that of all other imidazole elution samples of recombinant 6-His Stx2e A/B and STEC 150229 culture supernatant. The cytotoxicity of eluted sample with 500 mM imidazole was measured in 10^5^ TCCD_50_/100 µl at 48 h of incubation. The other samples showed the lower cytotoxicity, 10^3^ to 10^4^ TCCD_50_/100 µl at 48 h of incubation (Table 1).

## Discussion

In this study, we designed recombinant 6-His Stx2e A/B plasmids based on the shiga toxin sequence of STEC isolates to produce recombinant toxin proteins in a heterologous expression system. We also evaluated the cytotoxicity for the toxin protein complex active form of recombinant Stx2e A-His and intrinsic Stx2e B in STEC 150229. STEC 150229 showed that the production of shiga toxin was highest in a previous report [19]. From a total of 43 isolates from diagnosed case strains, 14 edema disease-causing *E. coli* strains including Stx2e and F18 genes were confirmed by toxin gene PCR. Among the edema disease-causing *E. coli*, 13 strains were observed to have cytotoxicity levels of 10 to 10^2^ TCCD_50_/100 µl, whereas STEC 150229 was observed to have a higher cytotoxicity level of 10^3^ TCCD_50_/100 µl in Vero cells [19] (data not shown). As it is known that these cells can synthesize and secrete the intrinsic shiga toxin to the extracellular region, we hypothesized that the overexpressed recombinant Stx2e A protein forms an active protein complex with the intrinsic Stx2e B component from STEC 150229. The recombinant Stx2e A contained the PelB peptide, the secretion leader, and it could be secreted into the periplasmic space of *E. coli*. Therefore, the Stx2e complex was collected and purified from the growth medium after protein overexpression. The cytotoxic activity of recombinant 6-His Stx2e A/B protein samples eluted with a higher imidazole concentration was significantly higher than that of all other imidazole elution samples. The cytotoxicity of recombinant 6-His Stx2e A/B from rSTEC 150229 samples eluted with 500 mM imidazole were measured in 10^5^ TCCD_50_/100 µl in Vero cells upon 48 h of incubation. Higher production of shiga toxin was obtained from recombinant 6-His Stx2e A/B. To detect purified recombinant Stx2e A and intrinsic B complex, we performed western blotting using mouse-generated anti-Stx2e serum. The rSTEC 150229 secreted higher concentrations of toxins than the wild-type STEC 150229 cells (Table 1). A nonspecific band detected in the negative control appeared to represent the protein originating from *E. coli* cells. However, purified Stx2e A protein was not detected. The detection limit of western blotting is known to be on the order of picograms depending on the ECL sensitivity. This indicates that our purified protein might have high toxicity even when present at only a low level.

## Conclusion

In conclusion, the recombinant Stx2e A protein forming an active protein complex with the intrinsic Stx2e B component from STEC 150229 and rSTEC 150229 produced high levels of shiga toxin.

## Acknowledgements

This research was supported by a grant from the Next-Generation BioGreen Program (PJ016225012021), Rural Development Administration, Republic of Korea, and by the National Research Foundation of Korea (NRF) grant funded by the Korea government (NRF-2020R1I1A2074647).

## Author’s contributions

**Conceptualization:** Jung Hee Park, Won Il Kim, Jin Hur

**Data curation:** Byoung Joo Seo, Jeong Hee Yu, Seung Chai Kim, Byung Yong Park

**Supervision:** Jung Hee Park, Won Il Kim, Jin Hur

**Writing-original draft:** Byoung Joo Seo, Jeong Hee Yu

**Writing-review & editing:** Jung Hee Park, Jin Hur

## Supporting information

**S1 Table.** Cytotoxic levels of recombinant Stx2e A-fragment and Stx2e B in Vero cell.

| Samples | Imidazole Conc.<br>(mM) | 10 fold dilutions |  |  |  |  |  |
| --- | --- | --- | --- | --- | --- | --- | --- |
|  |  | 1 | 2 | 3 | 4 | 5 | 6 |
| BL21 |  | -<sup>a)</sup> | - | - | - | - | - |
| 150229 |  | +<sup>b)</sup> | + | + | + | - | - |
| BL21 +<br>Stx2e A-fragment<br>(pStx2eA-fusion) | flow | - | - | - | - | - | - |
|  | 20 | - | - | - | - | - | - |
|  | 50 | - | - | - | - | - | - |
|  | 250 | - | - | - | - | - | - |
|  | 500 | - | - | - | - | - | - |
| BL21 +<br>Stx2e B<br>(pStx2eB) | flow | - | - | - | - | - | - |
|  | 20 | - | - | - | - | - | - |
|  | 50 | - | - | - | - | - | - |
|  | 250 | - | - | - | - | - | - |
|  | 500 | - | - | - | - | - | - |
| Elution buffer | 20 | - | - | - | - | - | - |
|  | 50 | - | - | - | - | - | - |
|  | 250 | - | - | - | - | - | - |
|  | 500 | - | - | - | - | - | - |
| Control |  | - | - | - | - | - | - |

<sup>a)</sup> None cytotoxicity; <sup>b)</sup> Cytotoxicity

**S1 Fig.**
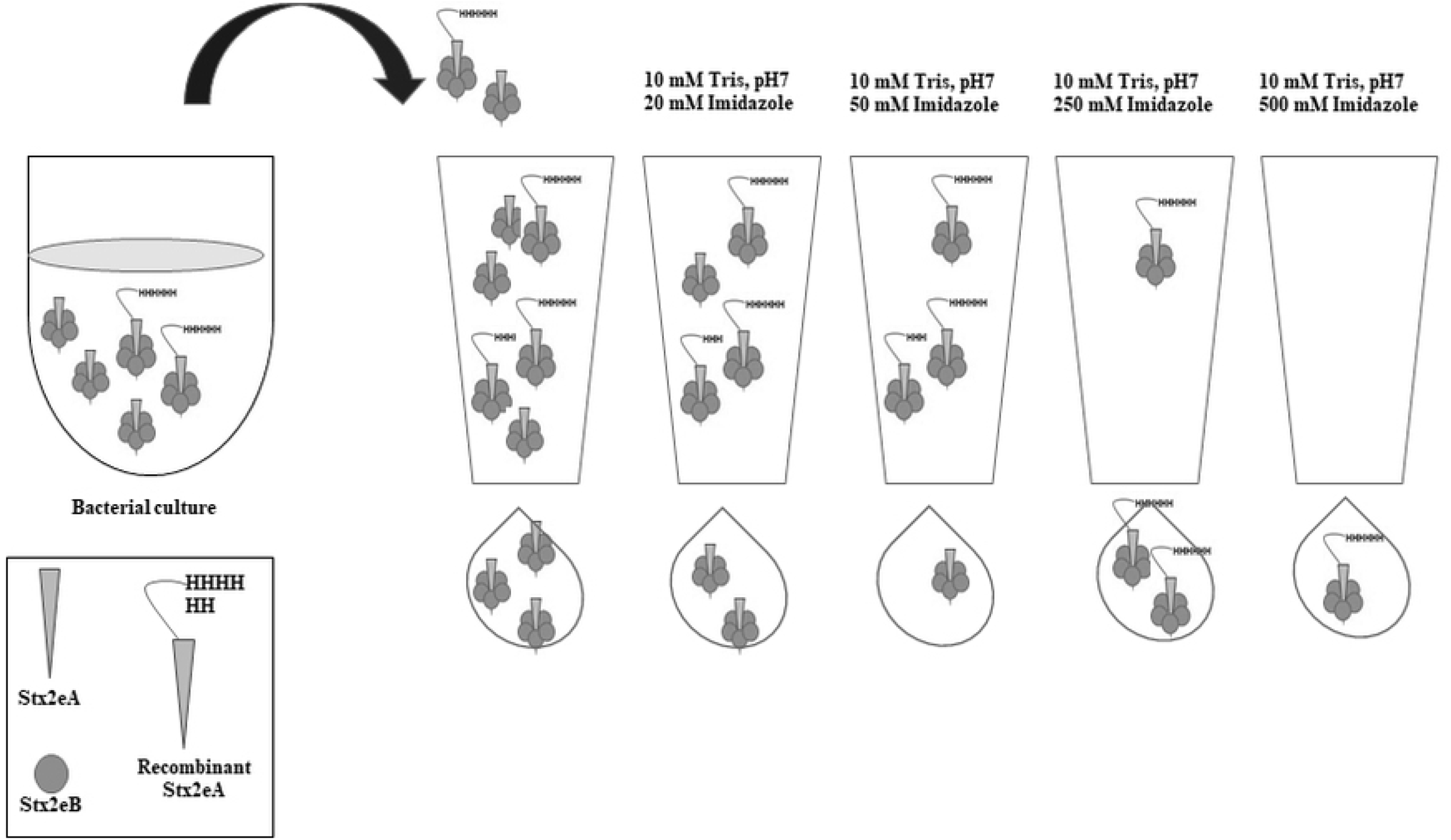
Scheme of the purification of recombinant Stx2e complex in STEC 150229 cells. The innate Stx2e complex was washed with a low imidazole concentration and the recombinant Stx2e complex was eluted in an increasing imidazole concentration.

